# A fence function for adherens junctions in epithelial cell polarity

**DOI:** 10.1101/605808

**Authors:** Mario Aguilar-Aragon, Alex Tournier, Barry J Thompson

**Author notes:** Corresponding author; @thompsonlab.

## Abstract

Adherens junctions are a defining feature of all epithelial cells, providing cell-cell adhesion and being essential for cell and tissue morphology. In *Drosophila*, adherens junctions are concentrated between the apical and basolateral plasma membrane domains, but whether they contribute to apical-basal polarisation itself has been unclear. Here we show that, in the absence of adherens junctions, apical-basal polarity determinants can still segregate into complementary domains, but control of apical versus basolateral domain size is lost. Manipulation of the level of apical or basal polarity determinants in experiments and in computer simulations suggests that junctions provide a moveable diffusion barrier, or fence, that restricts the diffusion of polarity determinants to enable precise domain size control. Movement of adherens junctions in response to mechanical forces during morphogenetic change thus enables spontaneous adjustment of apical versus basolateral domain size as an emergent property of the polarising system.

## Introduction

Cell polarity is a fundamental characteristic of living organisms. The molecular determinants of cell polarity have been revealed through pioneering genetic screens in yeast (*Saccharomyces cerevisiae* & *Schizosaccharomyces pombe*), worms (*Caenorhabditis elegans*), and fruit flies (*Drosophila melanogaster*) [1–5]. In yeast, the GTPase Cdc42 is essential for polarity [6] and has been shown to polarise via self-recruitment to the plasma membrane in a positive feedback loop [7–10]. In worms, Cdc42 acts with the PAR-3/PAR-6/aPKC complex at the anterior pole of the fertilised egg (zygote) to exclude the LGL/PAR-1/PAR-2 proteins to the posterior pole via mutual antagonism during mitosis [11–16]. Computer simulations of mutual antagonism in the worm zygote indicate that domain size is determined by the relative levels of anterior and posterior determinants [17]. Importantly, in the worm zygote, polarity determinants can diffuse freely across the domain boundary, which is not sharply defined but rather consists of overlapping gradients [18].

## Results and Discussion

In *Drosophila*, apical-basal polarisation of neural stem cells (neuroblasts) appears to be similar to the worm zygote, with Cdc42 acting apically with Par3/Bazooka (Baz), Par6 and aPKC during mitosis to exclude Lgl and other proteins basally via mutual angatonism [19–23]. As in the worm zygote, overlapping gradients of apical and basal determinants can be observed in neuroblasts and in computer simulations of apical-basal polarity (Fig 1A,B). In contrast, apical-basal polarisation of epithelial cells is different, with a sharp boundary forming between the same set of apical and basolateral determinants (Fig 1C). Furthermore, the relative size of the apical and basolateral domains of epithelial cells changes as cells transition from columnar to cuboidal or squamous cell shapes during tissue morphogenesis (Fig 1C-E). This observation suggests the existence of a diffusion barrier between the apical and basolateral domains of epithelial cells that can be repositioned by mechanical forces acting during development. In support of this view, addition of a diffusion barriers to computer simulations of apical-basal polarity [24] is sufficient to create a sharp boundary between apical and basal determinants and repositioning of the diffusion barriers is sufficient to determine apical versus basal domain size (Fig 1F-H).

**Figure 1.**
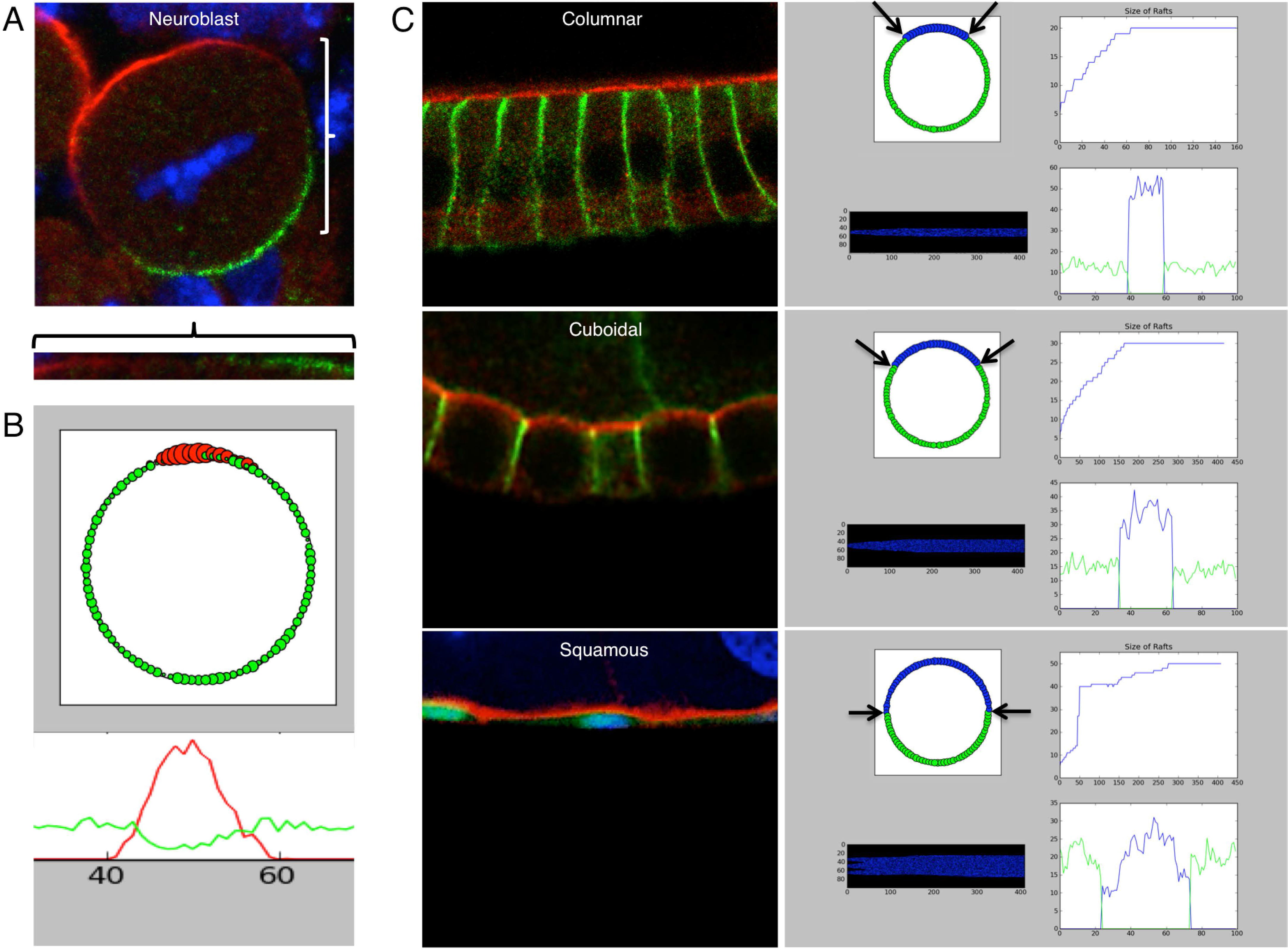
Addition of diffusion barriers to computer simulations of polarity enables sharp boundaries and domain size control. (A) *Drosophila* neuroblast stained for apical aPKC (red) and basal Miranda (green) to highlight overlapping gradients at the boundary between the two domains. (B) Computer simulation of apical (red) and basal (green) polarity determinants also generates overlapping gradients at the boundary between the two domains. (C) Columnar, cuboidal and squamous epithelial cells of the *Drosophila* ovarian follicular epithelium have sharply defined apical (red) and basolateral (green) domains, with the apical domain spontaenously adopting a very different size relative to the basolateral domain depending on cell shape. Computer simulations of cell polarity show that apical domain (blue) can change in size relative to the basal domain (green) spontaneously in response to the arbitrary shifting of diffusion barriers. Note also that the addition of diffusion barriers to the simulation creates a sharp border between apical and basal domains (compare with B).

The obvious candidate for such a diffusion barrier is the adherens junction, which is known to concentrate between the apical and basolateral domains in *Drosophila* epithelia but not neuroblasts [25–28]. While adherens junctions are known to be essential for epithelial cell shape [29–32], their exact role in polarisation of epithelial polarity determinants is still unclear. We therefore removed adherens junctions from the developing follicle cell epithelium of *Drosophila* by silencing of alpha-catenin expression and examined the localisation of the apical determinant aPKC and basolateral determinant Dlg (Fig 2A-D). We find that apical domain formation still occurs upon loss of alpha-catenin, despite the dramatic rounding up of cell shape and multilayering of cells (Fig 2D), consistent with previous results examining the loss of *beta-catenin/armadillo* in follicle cells [29]. Interestingly, we note that the size of the apical domain appears more variable and apical domain edges are no longer sharply defined in a ring when cells lack adherens junctions, unlike wild-type cells (Fig 2E,F). These findings indicate that adherens junctions are not required for apical domain formation but are required for precise domain size control in epithelial cells.

**Figure 2.**
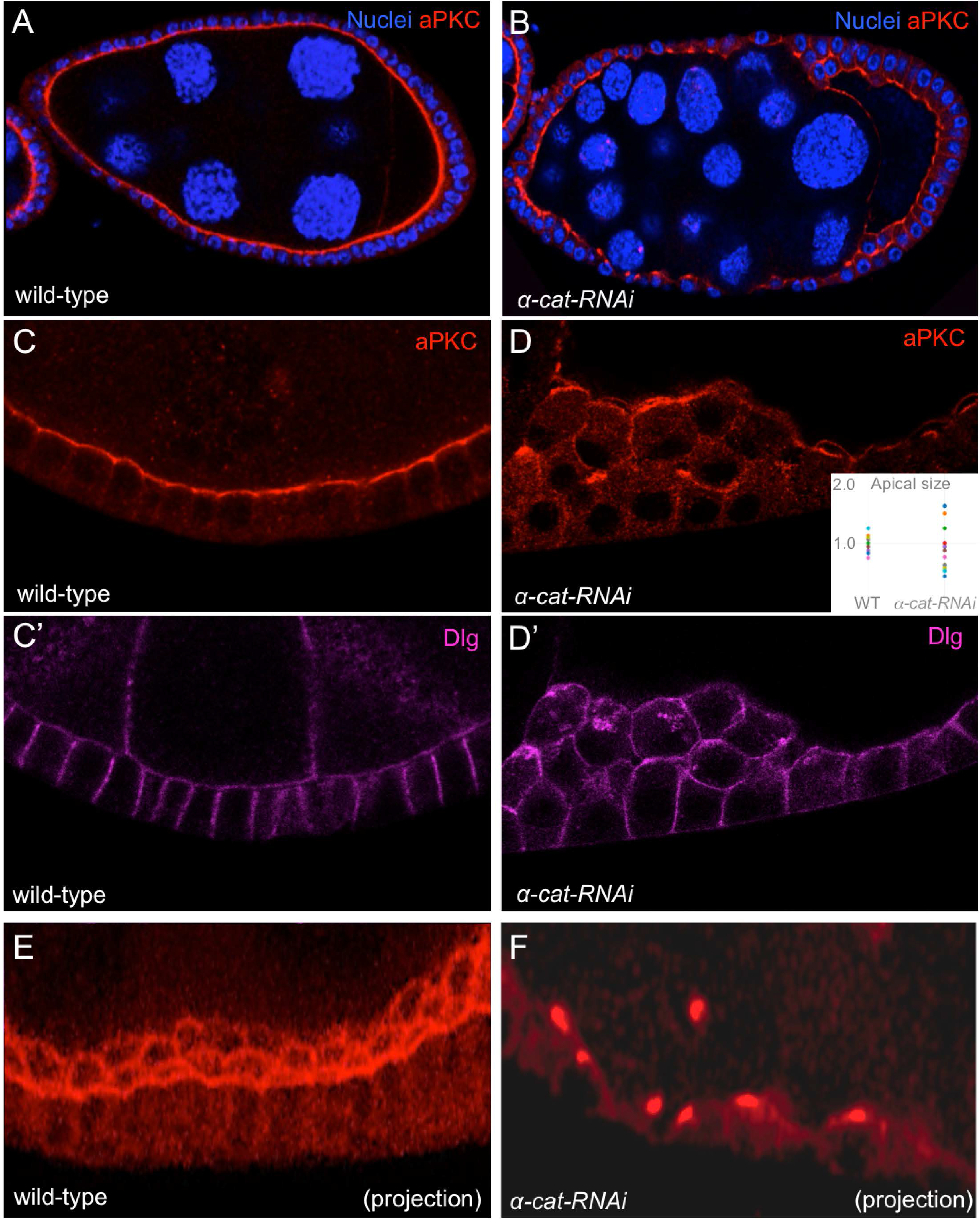
Adherens junctions are required for apical versus basolateral domain size control. (A) Wild-type *Drosophila* egg chamber stained for apical aPKC and nuclei. (B) Silencing of alpha-catenin by RNAi in follicle cells (TrafficJam.Gal4 UAS.alpha-catenin-RNAi) disrupts cell shape but still allows polarisation of aPKC. (C) Zoom of wild-type follicle cell epithelia stained for aPKC (red) and Dlg (purple; C’). (D) Zoom of alpha-catenin RNAi follicle cell epithelia stained for aPKC (red) and Dlg (purple; D’). Note that apical domain formation still occurs but that the size of the apical domain is abnormally variable (inset). (E) Projection of multiple sections of wild-type follicle cells stained for aPKC (red) showing concentration in an apical ring at the edge of the sharply defined apical domain. (F) Projection of multiple sections of alpha-cateninin RNAi follicle cells stained for aPKC (red) showing concentration in a single apical polar cap that is more variable in size than the wild-type.

We next tested whether adherens junctions act as diffusion barriers to limit the spread of apical or basolateral determinants in epithelia. We began by overexpressing the apical determinant Par3/Baz tagged with GFP in wild-type follicle cells and those lacking alpha-catenin (Fig 3A,B). We find that overexpressed Par3/Baz localises normally to the apical domain of wild-type epithelial cells, but spreads ectopically to form a larger apical domain in cells lacking alpha-catenin (Fig 3A-D). Similarly, overexpressing the basolateral determinant Lgl tagged with GFP has no effect on domain size in wild-type cells, but causes ectopic spreading of the basolateral domain in cells lacking adherens junctions (Fig 3E-H). These observations are paralleled in computer simulations of apical-basal polarity, where, in the absence of a diffusion barrier, increasing the levels of apical or basal determinants is sufficient to increase the corresponding domain size (Fig 3I,J). These findings show that adherens junctions function to restrict spreading of polarity determinants in epithelial cells, consistent with them acting as a diffusion barrier to control domain size.

**Figure 3.**
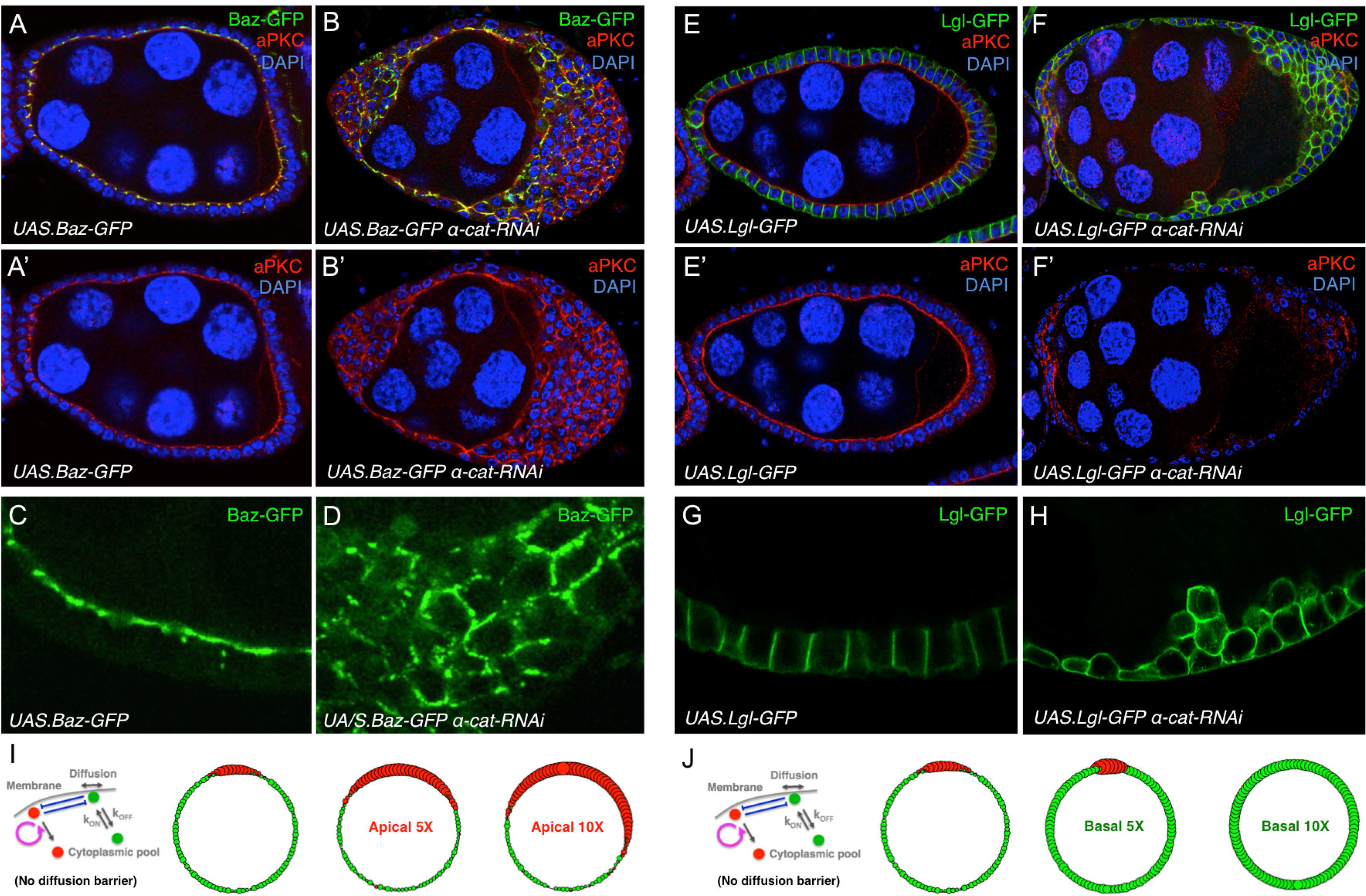
Adherens junctions provide a barrier that restricts spreading of apical and basolateral determinants. (A) *Drosophila* egg chamber overrexpressing UAS.Bazooka-GFP with the trafficjam.Gal4 driver. Stained for apical aPKC (red). DAPI marks nuclei (blue). (B) *Drosophila* egg chamber overrexpressing UAS.Bazooka-GFP and UAS.alpha-catenin RNAi with the trafficjam.Gal4 driver. Stained for apical aPKC (red). DAPI marks nuclei (blue). (C) Zoom of Bazooka-GFP overexpression in (A). Note normal apical localisation and domain size. (D) Zoom of Bazooka-GFP overexpression in alpha-catenin RNAi follicle cells in (B). Note spreading of Baz-GFP into a broader domain than in wild-type. (E) *Drosophila* egg chamber overrexpressing UAS.Lgl-GFP with the trafficjam.Gal4 driver. Stained for apical aPKC (red). DAPI marks nuclei (blue). (F) *Drosophila* egg chamber overrexpressing UAS.Lgl-GFP and UAS.alpha-catenin RNAi with the trafficjam.Gal4 driver. Stained for apical aPKC (red). DAPI marks nuclei (blue). (G) Zoom of Lgl-GFP overexpression in (E). Note normal basolateral localisation and domain size. (H) Zoom of Lgl-GFP overexpression in alpha-catenin RNAi follicle cells in (F). Note spreading of Lgl-GFP around the entire plasma membrane. (I) Computer simulation of polarised cells without diffusion barriers in which apical determinant levels are raised to drive apical spreading. (I) Computer simulation of polarised cells without diffusion barriers in which basal determinant levels are raised to drive basal spreading.

Finally, we re-visited the question of how adherens junctions come to be positioned between apical and basolateral polarity determinants. As expected, the ring of adherens junctions is located just below the apical domain and is disrupted upon loss of apical or basolateral determinants such as Cdc42 or Lgl (Fig 4A-C). We confirm that adherens junctions also require the key Rho-GTPase effector proteins Rho-kinase (Rok) and Diaphanous (Dia), which are known in mammalian cells to build the actomyosin contractile ring upon which junction formation depends [33–38], but whose role in *Drosophila* junction formation was recently called into question [39] (Fig 4D). Since Rho is localised uniformly at the plasma membrane, while its effector Rho-kinase (Rok) is localised apically (Fig 4E,F), we conclude that apical determinants must activate various RhoGEFs (Rho GTP Exchange Factors) specifically at the apical membrane, such that actomyosin then flows to lateral membranes where it can bind to E-cadherin at cell-cell contacts (Fig 4G,H), as also observed in live imaging of *Drosophila* embryos [40–42]. In support of the notion that positioning of adherens junctions requires apical actomyosin, sudden disruption of F-actin with Latrunculin A causes a failure of E-cadherin to concentrate in an apical ring, such that it redistributes along the entire lateral membrane (Fig 4I). In support of the notion that positioning of adherens junctions also requires actomyosin to flow to sites of cell-cell contact, overexpression of constitutive active RhoV14 is sufficient to induce actomyosin contractility around the entire plasma membrane, yet E-cadherin only accumulates ectopically in clusters at sites of cell-cell contact (Fig 4J). Thus, our findings support the model that apical basal polarity determinants induce actomyosin fibre networks apically so that adherens junctions then form at an apical-lateral position where E-cadherin is able to form homodimeric cell-cell contacts.

**Figure 4.**
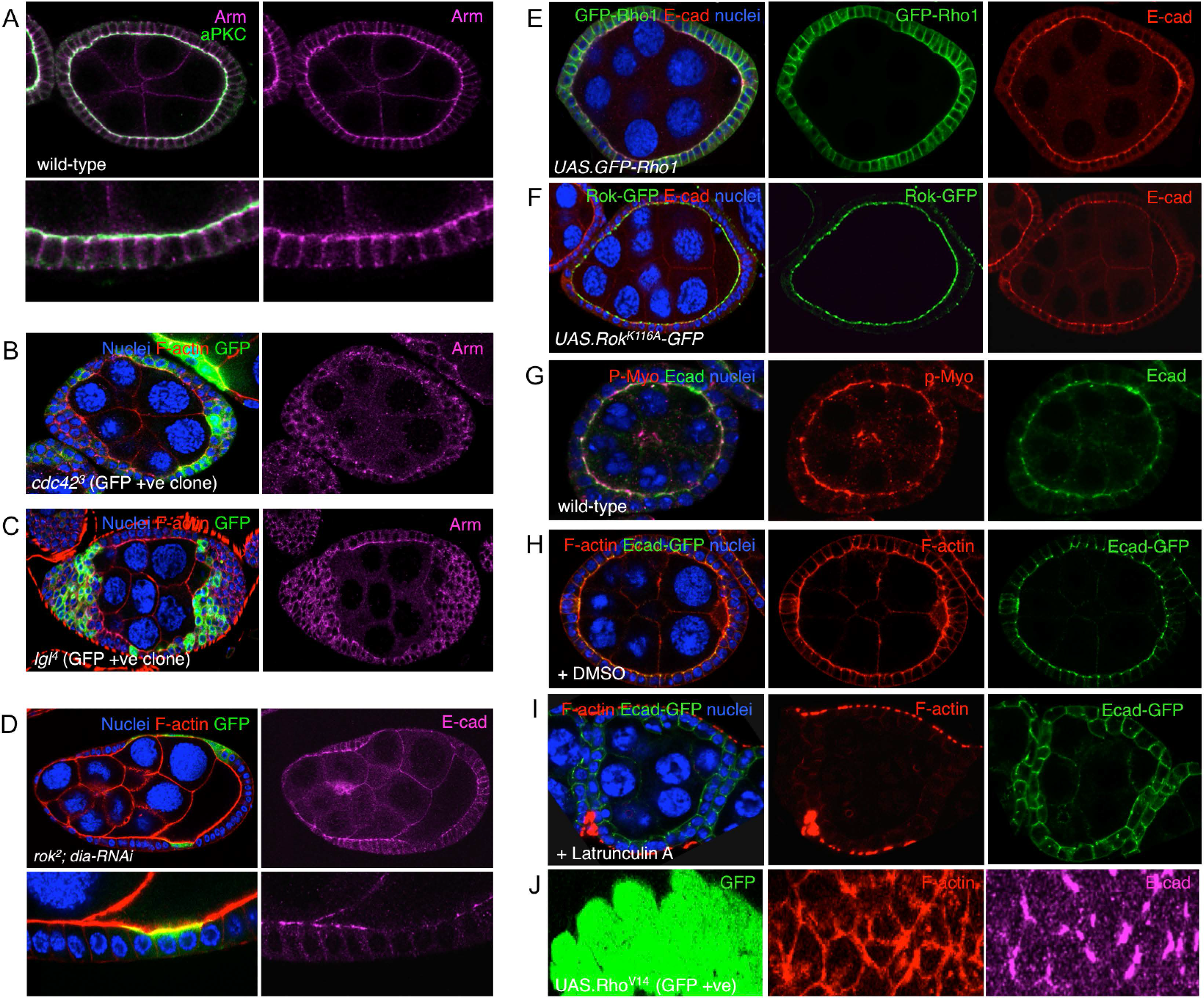
Apical-basal polarity determinants position the ring of adherens junctions via activating Rho-Rok-Dia to induce actomyosin apically. (A) Wild-type *Drosophila* egg chamber stained for apical aPKC (green) and junctional Armadillo/beta-catenin (purple). (B) *cdc42*^3^ mutant follicle cell clones, marked by the expression of GFP, lose the normal organisation of adherens junctions, marked by Armadillo/beta-catenin (purple). (C) *lgl*^4^ mutant follicle cell clones, marked by the expression of GFP, lose the normal organisation of adherens junctions, marked by Armadillo/beta-catenin (purple). (D) *rok*^2^ mutant follicle cells expressing UAS.Diaphanous-RNAi and UAS.GFP completely lose adherens junctions, marked by E-cadherin (purple). (E) Uniform localisation of GFP-tagged Rho1 expressed with trafficjam.Gal4 in the follicle cell epithelium. (F) Apical localisation of GFP-tagged Rho-kinase (Rok) expressed with trafficjam.Gal4 in the follicle cell epithelium. (G) Apical localisation of phospho-Myosin-II (p-Myo, red), a Rho-kinase substrate, and E-cadherin (Ecad, green) in the follicle cell epithelium. (H) Apical localisation of F-actin (red) and E-cadherin-GFP (Ecad-GFP, green) in the follicle cell epithelium (treated with DMSO as a control). (I) Loss of apical localisation of F-actin (red) and spreading of E-cadherin-GFP (Ecad-GFP, green) laterally in follicle cells treated with the actin depolymerising drug Latrunculin A. (J) Expression of constitutively active RhoV14 in clones of follicle cells marked by the expression of GFP (green), F-actin accumulates around the entire plasma membrane but E-cadherin only accumulates in clusters at cell-cell contacts.

In conclusion, cell polarity is more complex in epithelial tissues than in single-celled worm zygotes or neuroblasts. Epithelial cells use apical and basolateral determinants to position a ring of adherens junctions at the interface of the two domains via the classical mechanism of Rho GTPase mediated control of actomyosin. Once adherens junctions have formed, the boundary between apical and basolateral determinants is sharpened due to the action of adherens junctions as a diffusion barrier, or ‘molecular fence’. The importance of this fence function is revealed during tissue morphogenesis, when forces acting upon epithelial cells pull and push adherens junctions to drastically alter the shape of epithelial cells. The dynamic self-organising nature of apical-basal polarisation allows the relative size of each domain to spontaneously respond to repositioning of the adherens junction as cells change shape. This emergent property of the polarity system is illustrated by computer simulations of apical-basal polarity that can recapitulate this alteration of domain size in response to movement of diffusion barriers. Our findings indicate that changes in epithelial cell shape that expand or constrict the apical domain will therefore not require cells to program a corresponding alteration in the relative levels of apical-basal determinants, thus enabling rapid and dynamic tissue morphogenesis.

## Matherial and Methods

Expression of the UAS transgenic lines was achieved with either *tj.Gal4* and *GR1.Gal4* lines, and the actin ‘flip-out’ system. For ‘flip-out’ clones, 2 day old adult females were heat-shocked at 37°C for 1 hour and dissected 5 days after eclosion.

Mitotic clones were generated using the MARCM system and marked by the presence of GFP. In this case, third instar larvae were heat-shocked for 1 hour at 37°C and dissected 3 days after eclosion.

## Drosophila Genotypes

Fig 2A, C: *w*

Fig 2B, D: *w; tj.Gal4/+; UAS.α-cat-IR/+*

Fig 3A, C: *w; tj.Gal4/UAS.Baz-GFP+*

Fig 3B, D: *w; tj.Gal4/UAS.Baz-GFP+; UAS.α-cat-IR/+*

Fig 3E, G: *w; tj.Gal4/+; UAS.Lgl-GFP/+*

Fig 3F, H: *w; tj.Gal4/+; UAS.α-cat-IR UAS.Lgl-GFP/+*

Fig 4A,G,H,I: *w*

Fig 4B: *yw hs.flp tub.Gal80 FRT19A / cdc42^3^ FRT19A; tub.Gal4 UAS.GFP*

Fig 4C: *yw hs.flp tub.Gal4 UAS.GFPnls/+;lgl^4^FRT40A/FRT40A tub.Gal80*

Fig 4D: *yw hs.flp tub.Gal80 FRT19A / rok^2^ FRT19A; tub.Gal4 UAS.GFP;; UAS.dia-IR/+*

Fig 4E: *w UAS.GFP-Rho1/+;; GR1.Gal4/+*

Fig 4F: *w; UAS.venus-Rok^K116A^/+; GR1.Gal4/+*

Fig 4J: *w hs.flp; actin.FRT.STOP.FRT.Gal4 UAS.GFP; UAS.Rho^V14^*

### Immunostaining of ovaries and imaginal discs

Ovaries were dissected in PBS, fixed for 20 minutes in 4% paraformaldehyde in PBS, washed for 30 minutes in PBS/0.1% Triton X-100 (PBT) and blocked for 15 minutes in 5% normal goat serum/PBT (PBT/NGS). Primary antibodies were diluted in PBT/NGS and samples were incubated overnight at 4°C.

Secondary antibodies were used for 2 hours at room temperature and then mounted on slides in Vectashield (Vector Labs). Images were taken with a Leica SP5 confocal and processed with Adobe Photoshop.

Primary antibodies used were: rabbit anti-PKCζ (C-20) (1:250, Santa Cruz), mouse anti-Dlg (1:250, DSHB), mouse anti-Arm (1:100, DSHB), rat anti-E-Cad (DCAD2) (1:100, DSHB).

Secondary antibodies used were goat Alexa fluor 488, 546 or 647 (1:500, Invitrogen), Phalloidin (2,5:250, Life Technologies) to stain F-actin and DAPI (1μg/ml, Life Technologies) to visualize nuclei.

Wild type egg chambers were cultured in imaging media containing Schneider’s Media (Invitrogen), insulin (Sigma), heat-inactivated fetal calcium serum (GE Healthcare), trehalose (Sigma), adenosine deaminase (Roche), methoprene (Sigma) and Ecdyson (Sigma) with Latranculin A (0,05mM, Sigma), Cytochalasin D (0,05mM, Sigma) or DMSO (Sigma) as a control for 4 hours at room temperature. After treatment, samples were fixed and processed normally for imaging.

## Computational Model

Polarisation of a single cell was simulated using a stochastic modelling approach. The cell membrane was represented as 100 interconnected compartments, with the proteins (apical determinants ‘AD’ or basal determinants ‘BLD’) being able to diffuse from one compartment to the next. The cytoplasm was modelled as one ‘cytoplasmic pool’ compartment connected to all 100 membrane compartments. Proteins were able to interact with each other and had associated rates of coming on or off the membrane according to the rules and rates set out in the model (defined below). Using experimental constraints, the number of undefined parameters could be reduced to one, k_antag_, corresponding to the rate at which apical and basolateral determinants promote removal of one another from the membrane. Improving on the model by Altschuler et al 2008, we found that – for the modelling of polarity maintenance – the models could rely solely on the self-recruitment mechanism to maintain the apical determinants (AD) on the membrane. Thus the corresponding parameter, *k*_*on*_, which remained very small in Altschuler et al 2008, could be simply removed in the current model of polarity maintenance for AD.

### Stochastic event types

The five possible stochastic event types present in the model are:

- **Diffusion event**: a protein diffusing randomly from one compartment to a neighbouring compartment.
- **To cytoplasm events**: a protein randomly coming off the membrane.
- **From cytoplasm events**: a protein randomly coming onto the membrane from the cytoplasm.
- **Recruitment from cytoplasm**: a membrane bound AD randomly recruiting another AD protein from the pool to the membrane.
- **Membrane antagonism**: a membrane bound AD (or BLD) antagonising a membrane bound BLD (or AD), pushing it into the cytoplasmic pool.

### Assumptions and approximations

In the case where there is no BLD or no antagonism of BLD towards AD, the model reduces to that of Altschuler etal 2008. In this limit, there is no local antagonism and the system can be written using an Ordinary Differential Equation (ODE) following Altschuler etal, as:

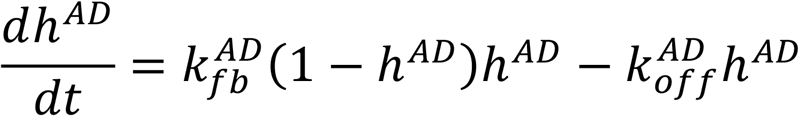

where *h*^*AD*^ is the proportion of ADs present on the membrane at any one time.

In steady state we get 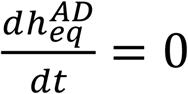, giving :

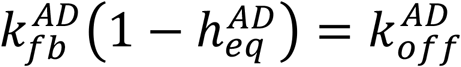

Given that we can measure 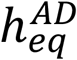 experimentally to be around 40% in the present biological system by quantitation of aPKC fluorescence intensity in imageJ, we have, 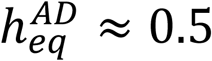, thus:

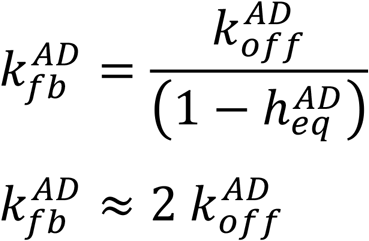

As soon as an antagonism between AD and BLD is introduced the equations cannot be simplified in the above manner and numerical methods have to be used. However, in practice, due to the fact that the overlap region between AD and BLD remains in most cases small, and hence the effect of this antagonism on the average cytoplasmic ratio remains small, this approximation was found to be useful in constraining the parameter space to the biologically meaningful values where 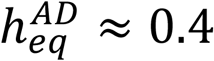.

In a similar fashion, we measured the membrane to cytoplasm ratio for Lgl-GFP with imageJ and we can solve the ODE for the case of BLDs in steady state without antagonism to ADs:

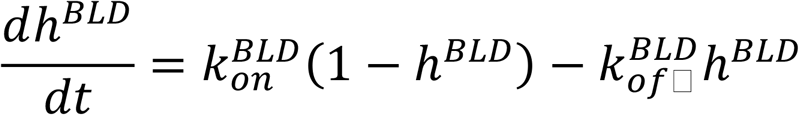

Which gives us for the steady state:

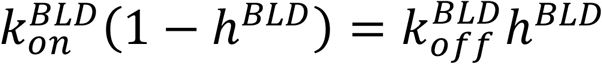

or

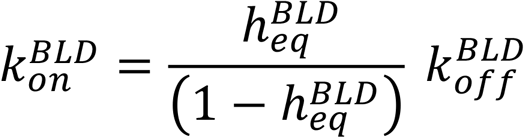

Which with measured to be 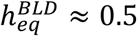 gives us:

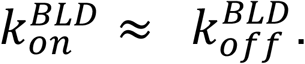

The rate at which ADs and BLDs spontaneously come off the membrane was assumed to be same for both species. The value for this parameter was estimated in Altschuler etal to be ≈ 9 min^−1^. Furthermore the rate at which ADs antagonise BLDs was assumed to be the same as the rate at which BLDs antagonise ADs. A satisfactory value for this number was found to be: *k*_*antag*_ = .1. A sensitivity analysis was performed with respect to this parameter (not shown).

### Final set of parameters

The following set of parameters was used:

| Parameter |  | Value | Evaluation |
| --- | --- | --- | --- |
| Diffusion of ADs along membrane<br>Diffusion of BLDs along membrane | $k_{diff}$ | $1.2 \mu\text{m}^2\text{min}^{-1}$ | From Altschuler etal 2008 |
| BLD from pool to membrane | $k_{on}^{BLD}$ | $\approx k_{off}^{AD}$ | Estimated using measured cytoplasmic ratio |
| AD from membrane to pool<br>BLD from membrane to pool | $k_{off}^{AD} = k_{off}^{BLD}$ | $9 \text{ min}^{-1}$ | From Altschuler etal 2008 |
| AD pulls AD from pool to membrane | $k_{fb}^{AD}$ | $\approx 2 k_{off}^{AD} = 18 \text{ min}^{-1}$ | Estimated using measured cytoplasmic ratio |
| AD pulls BLD off membrane<br>BLD pulls AD off membrane | $k_{antag}$ | $.1 \text{ min}^{-1}$ | Unknown |
| Number of ADs | $N_{AD}$ | 1000, 5000, 10000 | Variable parameter |
| Number of BLDs | $N_{BLD}$ | 1000, 5000, 10000 | Variable parameter |

### Algorithm and implementation

In each step of the algorithm an event type is first selected at random out of the set of possible event types according to probabilities dictated by the configuration of the system at that instant in time. Once the event type is selected, the compartment on which it will act is selected as random and the system updated according to that event. Finally, the time is updated by a small increment.

More precisely, the probability *P*_*i*_, of an event type, *i*, occurring at time, *t*, can be expressed as

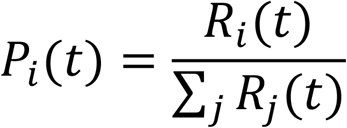

Where *R*_*j*_(*t*) is the overall rate of event type *j* at time *t*. For example, in the case where *i* corresponds to the event of having a BLD move from the cytoplasm to the membrane, the overall rate of that event is simply expressed as:

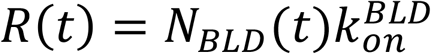

Rates corresponding to other event types can be calculated in similar fashion. Using the above expression an event type is randomly selected at each timestep of the simulation. In the case where the above BLD move is selected, a membrane compartment is randomly selected and a BLD moved from the cytoplasmic pool to that compartment. The time is then updated by a small increment *δt*:

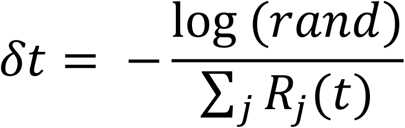

where *rand* is a random number between 0 and 1. The algorithm was implemented in Python using wxPython for the graphical interface and would typically run on a desktop PC in a few minutes.

